# Optimizing Neoadjuvant Treatment Response Prediction for Triple-Negative Breast Cancer Using Clinical Trial Data and Deep Auxiliary Learning

**DOI:** 10.1101/2024.11.18.620337

**Authors:** Chang In Moon, Bing Zhang

## Abstract

**Motivation:** Triple-negative breast cancer (TNBC) is an aggressive subtype of breast cancer with limited treatment options and poor prognosis. Developing predictive models for TNBC treatment responses is crucial but challenging due to data scarcity and the reliance on cell line data, which limits clinical translational value. Leveraging omics data from clinical trials, particularly through auxiliary learning, offers a potential solution to enhance predictive accuracy and reduce data requirements.

**Results:** In this study, we propose a new approach utilizing deep auxiliary task reweighting learning methods to automatically reweight auxiliary tasks, thereby optimizing the performance of the primary task of predicting TNBC treatment responses. We benchmark various auxiliary learning methods, including ARML, AdaLoss, GradNorm, and OL AUX, against traditional supervised machine learning algorithms and single-task learning baselines. Our results characterize the performance of auxiliary learning across various contexts, including utilizing parallel treatment arms within a multi-arm clinical trial, leveraging treatment arms from different clinical trials, and integrating multiple arms with the same treatment regimens across separate clinical trials. The last scenario also provides an opportunity for validating prediction models on an independent dataset, demonstrating the superior performance of the auxiliary learning models in predicting pathological complete response (pCR) in TNBC patients treated with standardized combinational chemotherapy with Taxane, Anthracycline, and Cyclophosphamide (TAC).

**Availability and Implementation:** Source code and additional resources can be accessed at https://github.com/moonchangin/DeepAux TxPred TNBC.

## Introduction

Breast cancer is a leading cause of cancer-related deaths worldwide (Lei et al., 2021; Sung et al., 2021). Triple-negative breast cancer (TNBC) is an aggressive subtype that accounts for approximately 15-20% of all breast cancers and is associated with poor prognosis and limited treatment options (Vagia et al., 2020; Lehmann et al., 2011). The previous standard of care for TNBC patients in the neoadjuvant setting was chemotherapy, with taxane and anthracycline/cyclophosphamide (TAC) forming the backbone of many chemotherapy regimens (Korde et al., 2021). The recent addition of immune checkpoint blockade has increased the response rate from 50% to 80%, but it comes with an increased risk of toxicity and high cost (Rizzo et al., 2022; Sternschuss et al., 2021). Identifying patients who can benefit from chemotherapy alone, and matching TNBC patients to the most appropriate chemotherapy remain important clinical challenges.

Molecular profiling of tumors by next-generation sequencing and microarray technologies is an area of intense investigation (Malone et al., 2020). This has led to an increase of genomic and transcriptomic characterization of TNBC tumors. Various datasets are publicly available with both transcriptomics data and treatment responses. Since these datasets profile tumors at baseline prior to treatment, association of molecular data with neoadjuvant response allows the identification of predictive biomarkers in patients likely to benefit from a specific treatment, thereby potentially optimizing treatment selection.

The typical approach to generate treatment response prediction models involves using pre-selected gene panels from a single study examining one treatment arm (Tabchy et al., 2010; Sayaman et al., 2020). The performance of these models is limited by many factors, with sample size being a major one. Unlike traditional methods that rely on only one, treatment-specific dataset/arm, our hypothesis centers on the potential of auxiliary learning, which utilizes auxiliary tasks across multiple TNBC clinical trials, to outperform established models in predicting pathologic complete response (pCR). pCR refers to the absence of any detectable cancer cells in tissue samples taken from the tumor site after neoadjuvant therapy, which is treatment given before the primary treatment (usually surgery). Achieving pCR is often associated with a better prognosis and longer survival in certain types of cancer, such as TNBC. On the other hand, non-pCR indicates the presence of residual cancer cells despite the treatment. Identifying biomarkers predictive of pCR can help tailor treatment strategies, improve patient outcomes, and reduce unnecessary exposure to potentially toxic therapies. By focusing on predicting pCR using molecular profiling and machine learning models, this study aims to enhance the precision of treatment selection and improve therapeutic outcomes for TNBC patients.

Auxiliary learning leverages additional tasks to improve the performance of the primary task by providing supplementary information that helps refine the model (Vafaeikia et al., 2020). This study explores whether integrating auxiliary tasks can improve predictive accuracy and reduce data requirements for TNBC treatment prediction. We utilize various deep auxiliary task reweighting learning methods to automatically reweight auxiliary tasks, thereby optimizing the main task’s performance. Our approach extends the application of auxiliary reweighting methods, traditionally used in the computer vision domain, to tabular data. This conceptual innovation not only improves the robustness of independent validation but also surpasses the performance of traditional machine learning methods and batch correction techniques. We compared auxiliary learning methods with traditional methods based on transcriptomics data from TNBC clinical trials, with treatment response information. Our benchmarking includes auxiliary learning methods Auxiliary Task Reweighting for Minimum-data Learning (ARML) (Shi et al., 2020), Adaptive Loss Balancing (AdaLoss) (Hu et al., 2019), Gradient Norm (GradNorm) (Chen et al., 2018), and Online Auxiliary Learning (OL AUX) (Lin et al., 2019), and compare them with several widely used traditional supervised machine learning algorithms.

## Material and Methods

### Data Sources

We collected transcriptomic data from 27 treatment arms across 16 TNBC clinical trials, involving a total of 1718 patients, all sourced from the ClinicalOmicsDB database (Moon et al., 2023). The primary source of transcriptomic data in ClinicalOmicsDB was from Gene Expression Omnibus (GEO). To pinpoint relevant datasets from clinical trial samples, specific search terms were used, including ‘cancer treatment response’, ‘neoadjuvant therapy’, ‘clinical trial’, and ‘clinical trial with treatment response’. These inclusion criteria resulted in clinical studies that included RNA profiling of treatment-naïve tumors and neoadjuvant treatment response labels. To ensure the accuracy of sample and treatment response information, we cross-referenced the original publications and corresponding data within https://clinicaltrials.gov/.

### Dataset Construction

Every treatment arm within a clinical trial was treated as an individual unit for dataset construction. Each of these datasets included gene expression data and clinical treatment response information of all patients encompassed within the respective treatment arm. Patients that achieved pCR were annotated as responders, all others were annotated as non-responders. Details on the gene expression and treatment response data processing are available in ClinicalOmicsDB paper (Moon et al., 2023).

### Deep Auxiliary Task Reweighting Learning Algorithm

To address the data scarcity problem in TNBC treatment prediction, we utilize deep auxiliary task reweighting learning methods. Our approach involves leveraging multiple auxiliary tasks to improve the main task’s performance, even with limited labeled data. The process is guided by the principle of reweighting auxiliary tasks to act as a surrogate prior for the main task, thus minimizing the divergence between the surrogate prior and the true prior.

#### Model Architecture

We assume a main task with training data *T*_*m*_ and *K* different auxiliary tasks with training data *T*_*ak*_ for the *k*-th task, where *k* = 1, …, *K*. The model includes a shared backbone parameterized by *θ*, with separate heads for each task. Our objective is to determine the optimal parameter *θ*^*^ for the main task by incorporating data from both the main and auxiliary tasks.

The training process involves minimizing the following joint loss function:

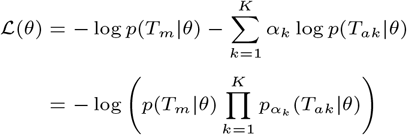

Here, *α* = (*α*_1_, …, *α*_*K*_) represents the set of task weights.

#### Objective Function

The core of our approach is based on the minimization of the Kullback-Leibler (KL) divergence between the true prior *p*^*^(*θ*) and a surrogate prior *p*_*α*_(*θ*), which combines the main and auxiliary task likelihoods:

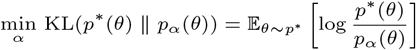

However, since *p*^*^(*θ*) is not accessible, we approximate it using the parameter distribution induced by the data likelihood of the main task *p*(*T*_*m*_|*θ*), denoted as *p*_*m*_(*θ*) ∝ *p*(*T*_*m*_|*θ*). This leads us to:

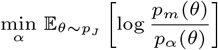

Here, 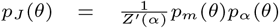, and *Z*^*′*^(*α*) is the normalization term. To estimate *p*_*J*_ efficiently, we use Langevin dynamics:

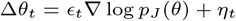

where *ϵ*_*t*_ is the learning rate, and *η*_*t*_ ∼ *N* (0, 2*ϵ*_*t*_) is Gaussian noise.

#### Optimization

Our optimization objective is refined to minimize the Fisher divergence between the score functions of the main task and the surrogate prior:

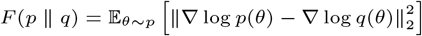

Thus, our final objective becomes:

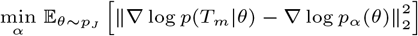

where ∇ log *p*_*m*_(*θ*) = ∇ log *p*(*T*_*m*_|*θ*).

#### Algorithm

##### Algorithm 1

Auxiliary Task Reweighting for Minimum-data Learning

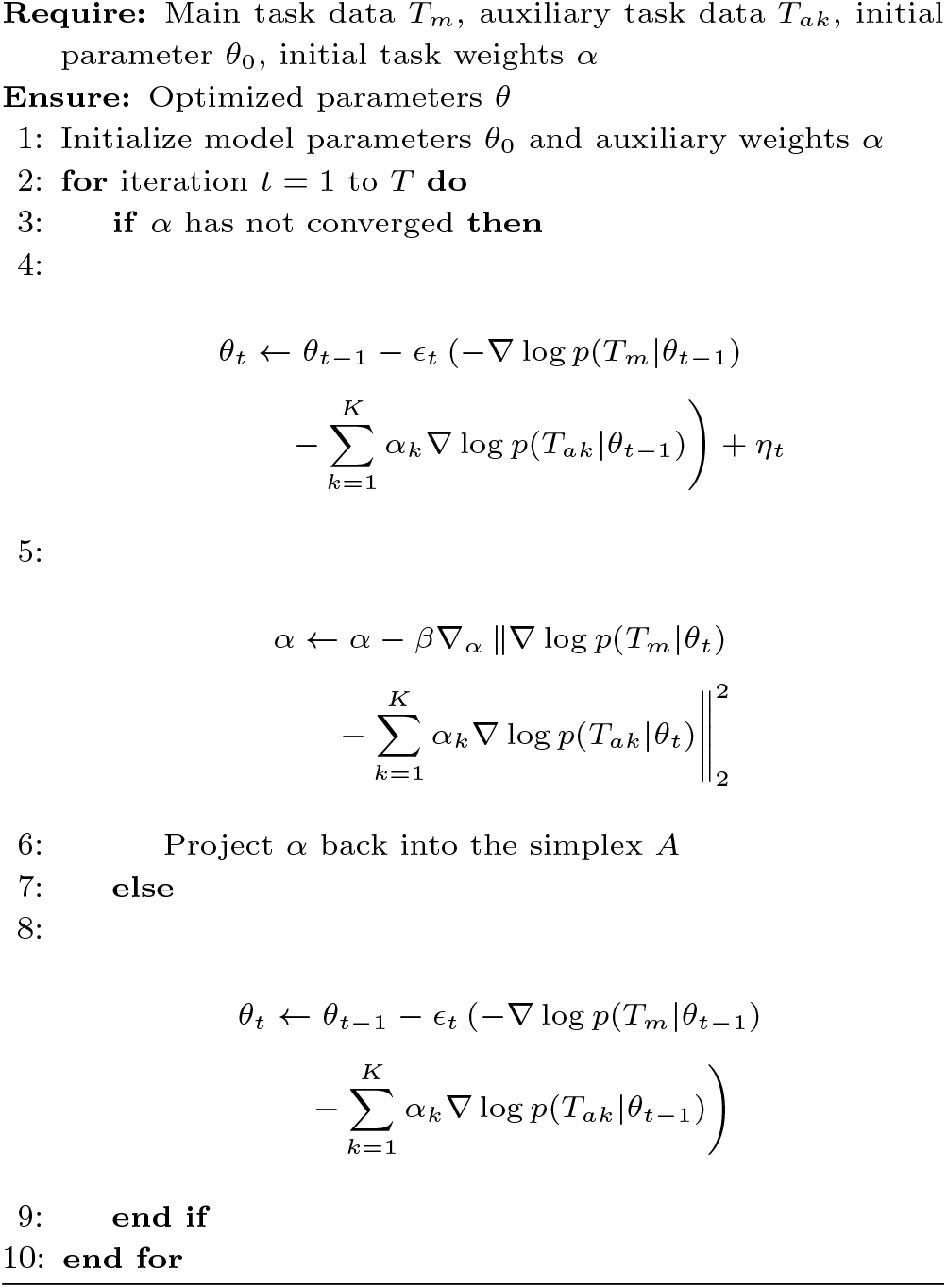

Initial values of the model parameters *θ* and task weights *α* are initialized randomly under a Gaussian distribution.

This random initialization helps avoid any inherent bias in the starting point of the optimization process and allows the gradient-based optimization methods to explore a broad range of possible solutions. The algorithm iterates until the task weights *α* have converged. Convergence is defined when the change in task weights *α* between consecutive iterations falls below a predefined threshold (e.g., *ϵ* = 10^−5^). During this process, both the task weights and model parameters *θ* are updated iteratively to ensure optimal performance. Regular gradient descent is applied to update the model parameters once convergence of *α* is achieved. To prevent overfitting, early stopping is employed if no significant improvement in validation loss is observed over a set number of epochs (e.g., 10 epochs) or if a maximum number of iterations (e.g., 100 epochs) is reached. The multilayer perceptron model predicts labels for each task, but only the main task’s label is used for evaluation.

#### Auxiliary Learning Methods

We utilize various auxiliary learning methods, including ARML, AdaLoss, GradNorm, and OL AUX, as well as multiple widely used supervised learning algorithms, including Extreme Gradient Boosting (XGBoost) (Chen and Guestrin, 2016), Random Forest (RF) (Breiman, 2001), logistic regression (Cox, 1958), and single-task learning (STL), specifically referring to multilayer perceptrons (Haykin, 1994) in this study, for comprehensive evaluation. The scikit-learn package was used for these supervised learning methods and detailed hyperparameters are available in the GitHub repo.

In multi-task learning, accurately estimating the relationship between different tasks is crucial for balancing multiple losses. This process often involves task clustering via a mixture prior, which effectively screens out unrelated tasks but does not further differentiate among related tasks. Auxiliary task reweighting methods, including ARML, AdaLoss, GradNorm, and OL AUX, address this by assigning greater weights to auxiliary tasks that provide a more relevant representation of the main task. Specifically, the ARML algorithm reduces the data requirements of a primary task by adaptively reweighting auxiliary tasks during joint training. This algorithm uses the parameter distribution, shaped by the likelihood of auxiliary tasks, as a surrogate prior for the main task. It adjusts the task weights to minimize the divergence between the surrogate and true priors of the main task, thereby decreasing data requirements. Similarly, OL AUX estimates task relationships based on the similarity of gradients and employs the inner product as a similarity metric to expedite training. Another strategy involves managing multiple losses through techniques like the gradient norm (GRADNORM), which focuses on equalizing the influence of each task by adjusting gradients, or Adaptive Loss Balancing (ADALOSS), which addresses uncertainty by dynamically altering loss weights based on the learning state.

#### Batch Effect Removal

When multiple arms across separate clinical trials use the same treatment regimens, batch effect removal techniques can be applied to facilitate the integration of data from different trials in order to increase sample size for the training of traditional machine learning models. We employed two batch correction techniques widely utilized in the field. Specifically, z-score normalization and ComBat (Zhang et al., 2020) were applied to RNA-Seq data before model training and testing to reduce technical variability between datasets.

Z-score normalization standardizes each gene’s expression values by calculating the mean and standard deviation within individual datasets to be integrated, and then adjusting the values accordingly (subtracting the mean and dividing by the standard deviation) concatenating them into a single matrix. The resulting normalized data ensured that expression levels were comparable across different datasets. The same process was applied to the testing set for evaluating model performance. ComBat uses an empirical Bayesian framework to estimate and adjust batch-specific effects, accounting for both biological and technical variability. The batch effect correction was performed on the training datasets, and the learned parameters were then applied to the testing dataset. This ensured that the testing set underwent the same adjustments as the training data.

## Results

### Overall Workflow

**Figure 1** illustrates the workflow of the auxiliary learning framework used to predict treatment response. The main task involves predicting treatment response using data from the treatment arm of primary interest, while the auxiliary tasks use data from other presumably relevant treatment arms to support the main task. The main task dataset is split into two parts: 70% for training and 30% for testing. The training data is further divided into two subsets: 50% is used together with the auxiliary task datasets to train the main and auxiliary tasks, while the remaining 50% is allocated for validation. To reduce dimensionality and ensure feature consistency across datasets, the top 500 genes are selected using univariate analysis from 70% of training data. These genes are then transformed into 10 PCA components, which are used as input features for both the main and auxiliary tasks. During each epoch of training, the task weight (*α*) is adjusted, and the parameter (*θ*) is updated based on the convergence criteria. The model’s performance is evaluated on the validation set at each epoch. If the validation loss does not improve for 10 consecutive epochs, or if 100 epochs are completed, an early exit condition is triggered to prevent overfitting. The final model performance is evaluated using the area under the receiver operating characteristic curve (AUROC) on the reserved testing dataset. The final performance is evaluated using AUROC on the testing data, with results averaged over 30 Monte Carlo cross-validation (MCCV) iterations. The same workflow for cross-validation and feature selection is applied to supervised learning methods without using data from auxiliary treatment arms.

**Fig. 1.**
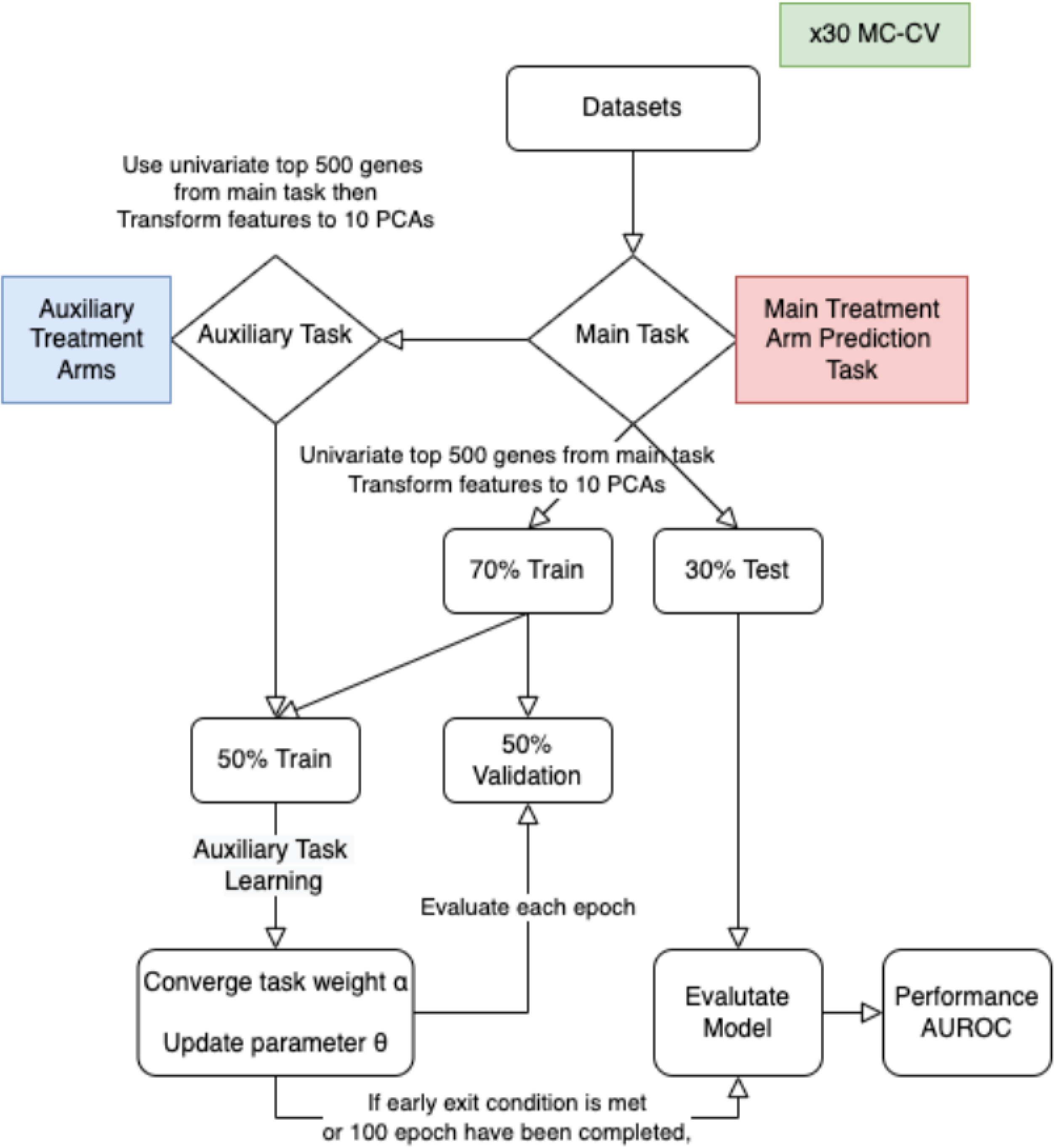
Auxiliary learning high-level model diagram

### Assessing Auxiliary Task Reweighting Through Simulation

Data from the BrightTNess trial was used to evaluate the efficacy of ARML, a representative auxiliary task reweighting algorithm, in predicting treatment responses. The study cohort included 482 patients, all of whom received anthracycline and cyclophosphamide backbone treatment after receiving the agents provided in one of the following three experimental arms:

- Arm A: Veliparib + Carboplatin + Paclitaxel (237 patients)
- Arm B: Carboplatin + Paclitaxel (122 patients)
- Arm C: Paclitaxel (123 patients)

The response rates (respond vs. not respond) for the treatment arms were 54:46, 57:43, and 33:67, respectively **(Figure 2A)**. For the primary task, we aimed to predict treatment responses of the Arm A regimens. To evaluate the impact of auxiliary tasks and determine the effectiveness of auxiliary task reweighting, we created the following datasets:

**Fig. 2.**
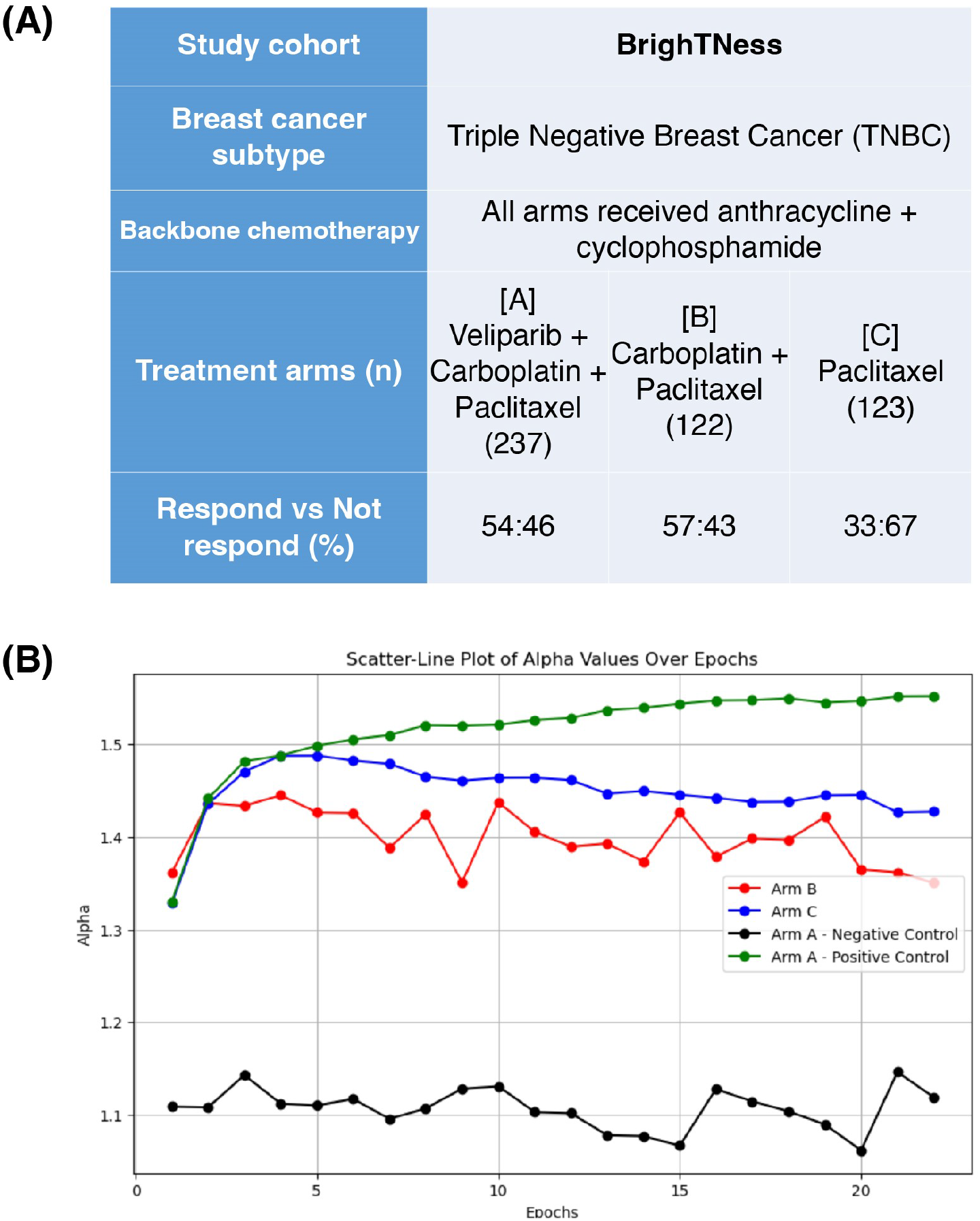
**(A)** Brightness Trial Setup and Evaluation Details: The table outlines the clinical trial study, breast cancer subtype, uniform treatment, treatment arms, and response rates. The training involved random selection of 100 samples from Arm A and creation of a pseudo auxiliary learning sample from 100 samples from Arm A. The evaluation was performed on the remaining 37 samples from Arm A. **(B)** Scatter-Line Plot of Alpha Values Over Epochs: Comparison of alpha reweighting for different auxiliary tasks in the Brightness Trial. The plot shows alpha values over epochs for four arms: Arm B (Red), Arm C (Blue), negative control with 100 samples from Arm A (Black), and positive control with 100 samples from Arm A (Green). The positive control represents the true prior (Arm A) before shuffling labels. The negative control represents Arm A with shuffled data and labels, which serves as a control to demonstrate irrelevance to the prediction task.

#### Main Task Dataset

- 100 samples from Arm A were randomly selected to form the main task dataset

#### Auxiliary Learning Datasets

- Positive Control: 100 samples from the remaining Arm A samples were randomly selected to form a pseudo-auxiliary learning task (i.e., representing the true prior).
- Negative Control: data and labels of the 100 samples from the positive control dataset were shuffled to simulate a non-informative auxiliary task.
- Auxiliary Task from Arm B: 122 samples from Arm B to assess cross-arm relevance.
- Auxiliary Task from Arm C: 123 samples from Arm C to assess cross-arm relevance.

#### Evaluation Dataset

- The evaluation was performed on the remaining 37 samples from Arm A.

**Figure 2B** presents the scatter-line plot of alpha values over epochs, comparing the alpha reweighting for different auxiliary tasks described above. The positive control auxiliary task (green line) consistently achieved higher alpha values compared to the other auxiliary tasks. In contrast, the alpha values for the negative control (black line) remained low and stable. These results demonstrate that the learning algorithm effectively distinguishes between auxiliary tasks that are relevant and those that are irrelevant to the main task, appropriately assigning weights to each task. Notably, the alpha values for the auxiliary tasks from Arm B (red line) and Arm C (blue line) were lower than those of the positive control but substantially higher than those of the negative control. This suggests that these parallel arms contribute valuable information to the prediction task from Arm A, likely due to shared chemotherapy agents or similar drug mechanisms across different chemotherapy agents.

### Performance of Auxiliary Learning Methods in a Multi-Treatment Arm Clinical Trial

We next evaluated the performance of various auxiliary learning methods in the multi-treatment arm clinical trial setting, where one treatment arm was used as the primary task while the others from the same trial served as auxiliary tasks. Parallel treatment arms within the same trial often share similar treatment regimens, which enhances the relevance between the main and auxiliary tasks. Additionally, patients are typically randomly assigned to different arms, and omics data are generated using consistent protocols, thereby minimizing potential batch effects. We continued using the BrighTNess trial in this analysis, rotating each of the three arms as the primary task while using the remaining two as auxiliary tasks for auxiliary learning. For traditional supervised learning, only the data from the primary task was used. **Figure 3** presents the comparative performance of four auxiliary learning methods and four supervised learning methods, based on results from 30 MCCVs. Auxiliary learning methods generally achieved significantly higher mean AUROC values compared to all four supervised learning methods across the three treatment arms. The performance gain was particularly evident for Arms B and C (**Figure 3c, e**), which had smaller sample sizes. In Arm A, which had the largest sample size, although AdaLoss showed inferior performance in some comparisons, the other three methods still significantly outperformed the supervised learning methods. These results demonstrate the potential of auxiliary learning methods to enhance prediction accuracy in the multi-arm clinical trial setting, especially when data is limited in a specific arm, while also maintaining strong performance with larger datasets.

**Fig. 3.**
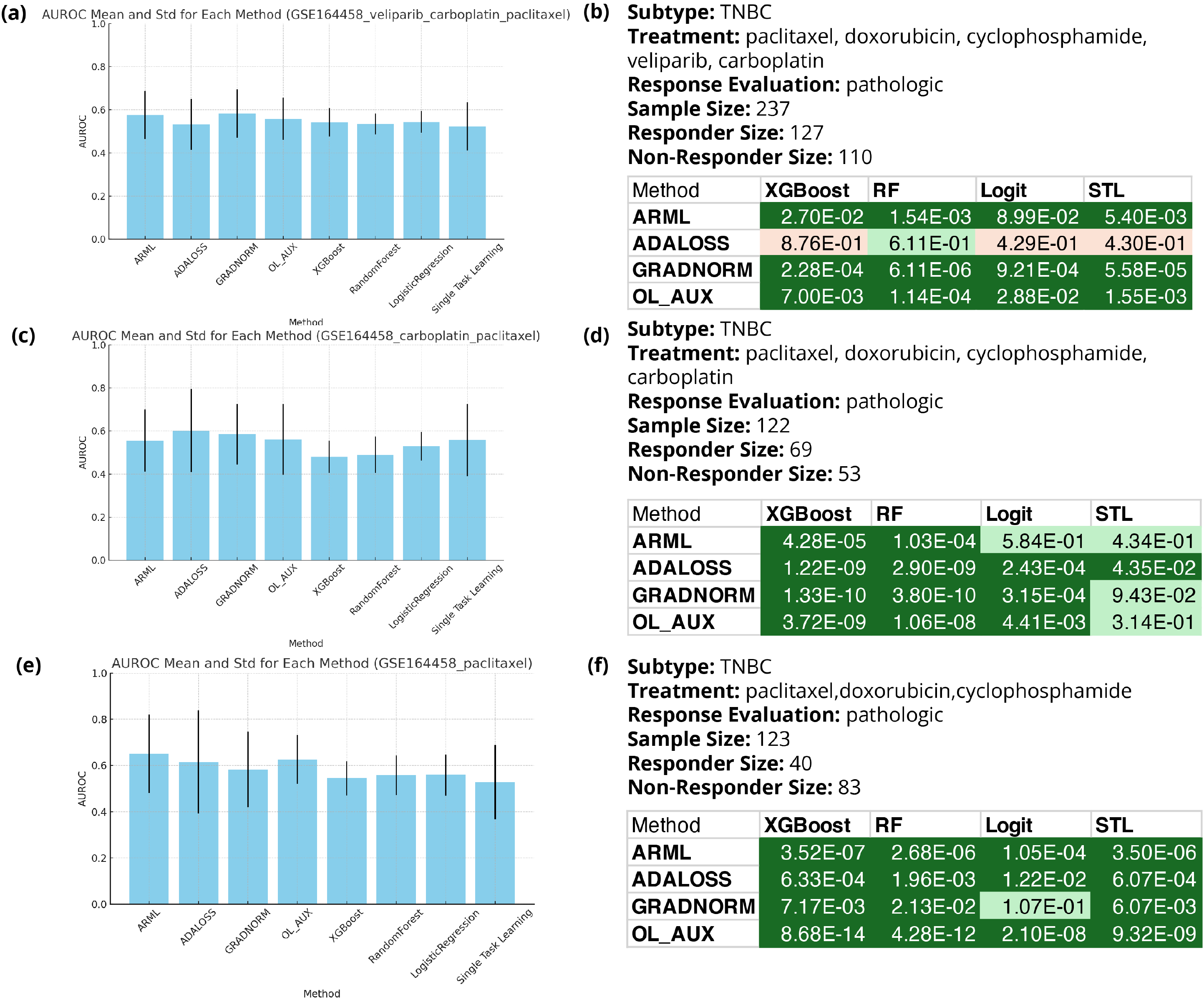
Comparison of deep auxiliary learning methods (ARML, ADALOSS, GRADNORM, OL AUX) with baseline models (XGBoost, Random Forest, Logistic Regression) and single-task learning (STL) in the BrighTNess cohort (GSE164458). Bar plots **(a, c, e)** show AUROC means and standard deviations from 30 MCCVs across three treatment arms. Tables **(b, d, f)** provide paired t-test results, with green indicating auxiliary methods outperforming baselines, orange showing the reverse, and darker colors representing statistically significant differences (p-value ¡ 0.05).

### Application of Auxiliary Learning to 27 TNBC Treatment Arms

Next, we expanded our evaluation to a multi-trial setting, using each of the 27 treatment arms across 16 TNBC clinical trials as the main task, with the remaining 26 serving as auxiliary tasks.

**Figure 4** presents a comparison of baseline supervised learning and single-task learning models to various deep auxiliary learning methods. The evaluation metrics are expressed as the best AUROC obtained from baseline models (XGBoost, RandomForest, Logistic Regression, STL) compared to auxiliary learning methods (ARML, ADALOSS, GRADNORM, OL AUX). Each study is plotted based on its best baseline and auxiliary learning performance across 30 MCCVs, with a 1:1 reference line shown to indicate equal performance between the two approaches. The dotted lines represent a ±0.1 AUROC range from the 1:1 line, indicating cases where the two methods yield similar results (within 0.1 AUROC). This range helps highlight cases where the differences in performance between baseline and auxiliary learning are minimal, making it easier to focus on datasets where auxiliary learning demonstrates more significant improvements or reductions in performance.

**Fig. 4.**
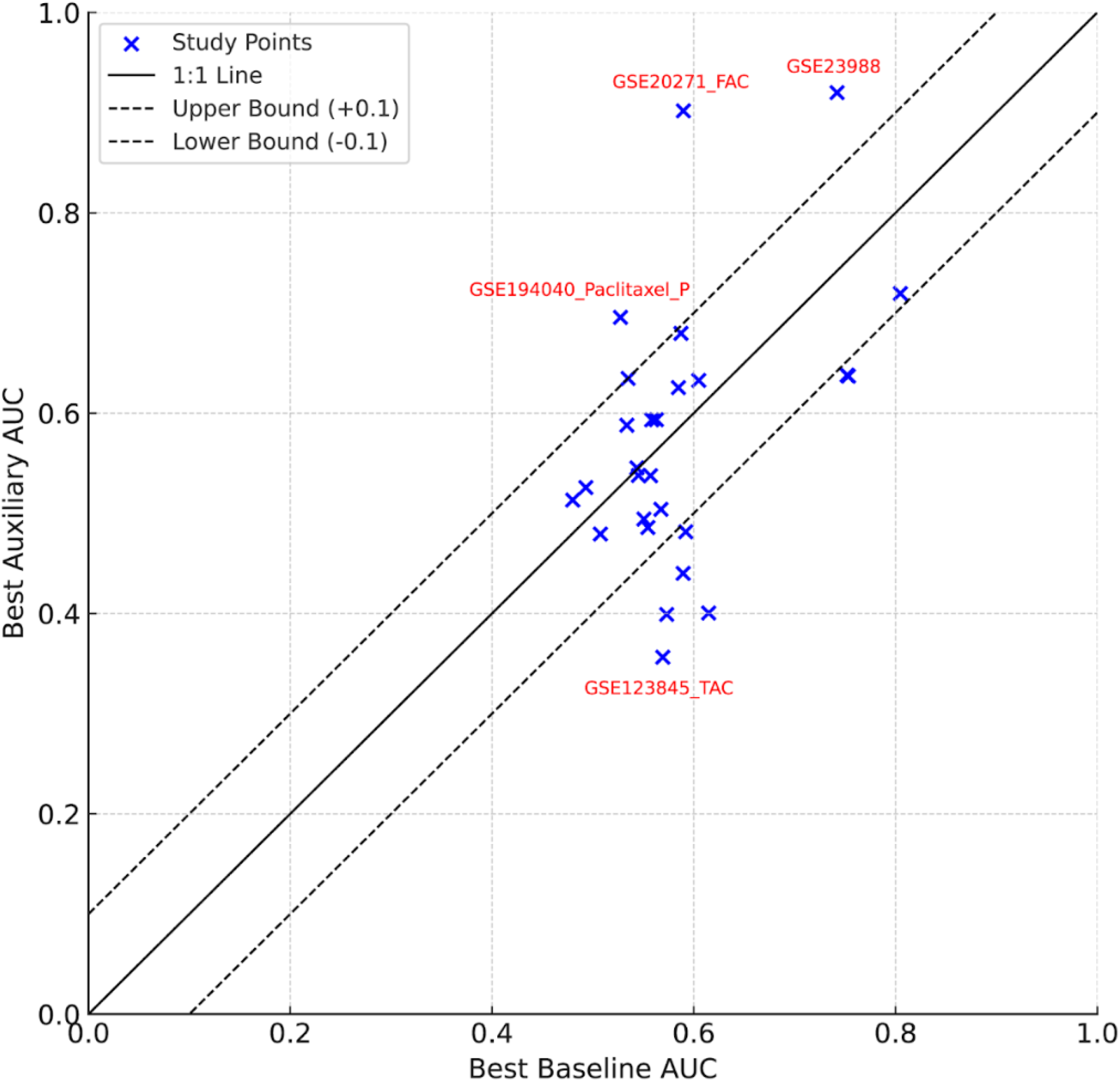
Scatterplot comparing the best baseline MCCV mean AUC performance and the best auxiliary learning AUC performance across various studies. The X-axis represents the best AUC obtained using baseline models (XGBoost, RandomForest, Logistic Regression, STL), and the Y-axis represents the best AUC from auxiliary learning methods (ARML, ADALOSS, GRADNORM, OL AUX). Each point corresponds to a study, and the black solid line represents the 1:1 line where baseline and auxiliary performances are equal. Dotted lines show the ±0.1 range around the 1:1 line. Studies where the auxiliary learning performance outperforms the baseline are labeled in red. Studies within the ±0.1 range of the 1:1 line (upper and lower bounds) are excluded from the labeling. This highlights the outlier studies where auxiliary learning shows a significant improvement or reduction in AUC compared to baseline models. Abbreviations: T=Taxane, A=anthracycline, C=cyclophosphamide, P=platinum, Pembro=pembrolizmab, F = fluorouracil.

While the two methods showed comparable performance on most datasets, as seen by the close clustering of points around the 1:1 line, auxiliary learning methods consistently achieved higher AUROC scores compared to baseline models in three datasets, including GSE20271 FAC (0.902), GSE23988 FAC (0.92), and GSE194040 Paclitaxel Pembrolizumab (0.691). For instance, in GSE194040 Paclitaxel Pembrolizumab, ARML significantly outperformed traditional machine learning models. GRADNORM also showed exceptional performance, particularly in GSE20271 FAC, achieving the highest AUROC value of 0.902. Comprehensive median 30 MCCV AUROCs and consistent strong performance across all auxiliary learning methods for these datasets are detailed in **Supplementary Table 1**.

A key factor contributing to these higher AUROC scores, especially in GSE23988 FAC and GSE20271 FAC, lies in the nature of the datasets themselves. GSE23988 FAC originates from a multi-national cohort, while GSE20271 FAC comes from a U.S.-based cohort. Both datasets use microarray platforms with the same treatment regimen, which may explain the superior performance of auxiliary learning methods. Specifically, these results suggest that auxiliary learning techniques are highly effective in identifying and leveraging information from closely related auxiliary datasets without prespecification, leading to improved performance. Thus, although immediate gains vary, the auxiliary learning approach provides a reliable framework for efficiently borrowing information from auxiliary datasets when possible, while maintaining comparable performance in other scenarios.

### Independent Validation Benchmarking on TAC Combination Therapy

The TAC regimen, frequently used as a control in clinical trials before PD-1 inhibitors were approved, provides a special scenario for aggregating datasets to increase sample size and potentially improve model performance for traditional supervised machine learning models. This also provides an opportunity for model validation using independent data, without relying on cross-validation. However, merging data from different trials is complex due to differences in sequencing platforms and batch effects, which make batch effect correction a common practice in supervised learning. We hypothesized that auxiliary learning, by assigning higher weights to similar datasets, could serve as a more effective data integration strategy than batch correction methods like z-score normalization or ComBat (Zhang et al., 2020).

**Figure 5A** displays the datasets used in this analysis, which focus on the TAC regimen across five clinical trials. GSE164458, with the largest sample size, was used for the main task in auxiliary learning, while GSE163882, with the second largest sample size, served as an independent validation dataset. Three additional datasets (GSE194040, GSE41998, and GSE123845) were employed as auxiliary datasets. For traditional machine learning, GSE163882 was used as the independent test data, while the other four were used for training. We first compared supervised learning using batch-corrected data versus uncorrected data. We observed minor improvements or even degraded performance of models using batch-corrected data **(Table 4-1, upper section)**.

**Fig. 5.**
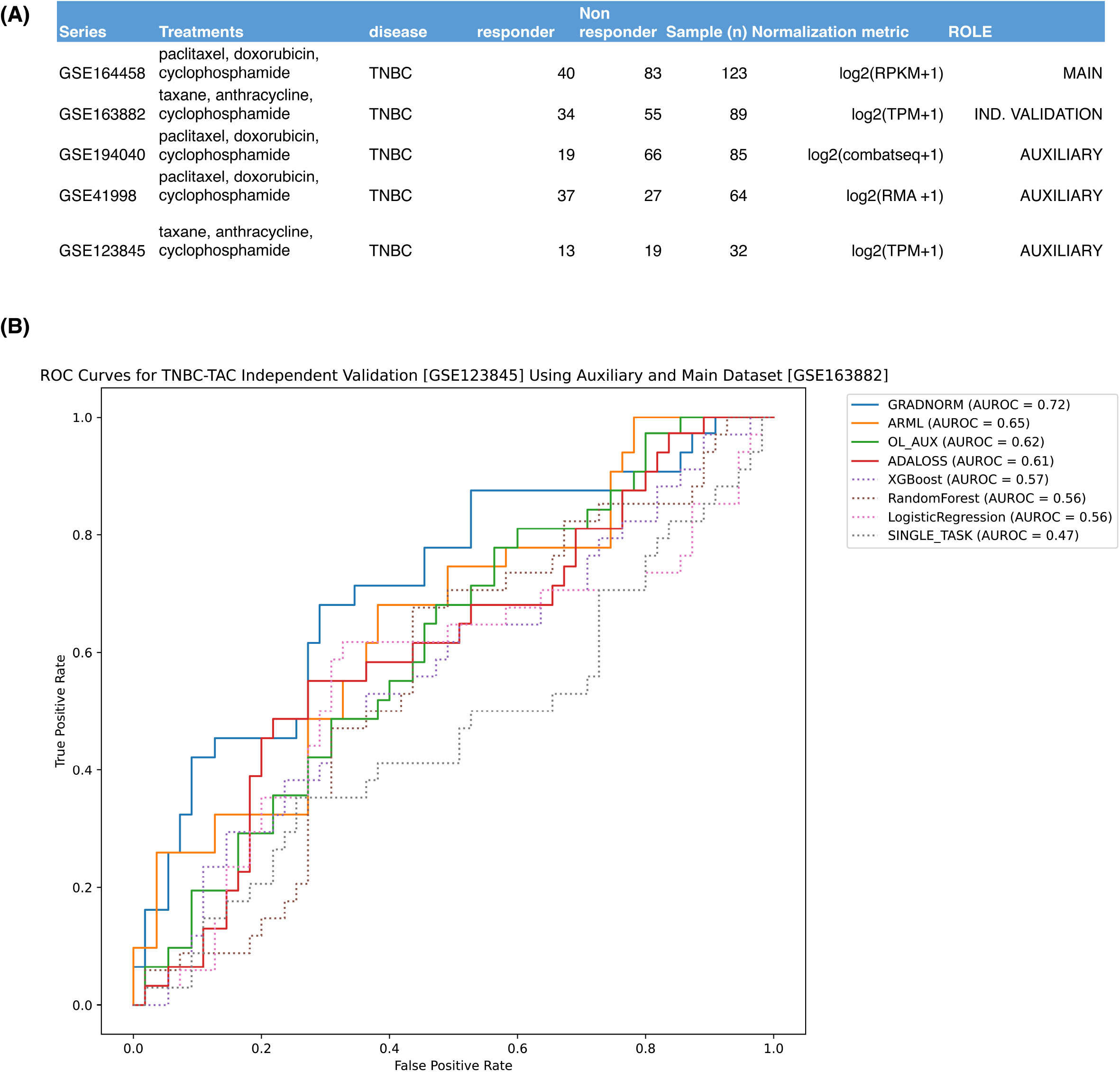
**(A)** Summary table of datasets used in the analysis, detailing the series, treatments, disease, number of responders and non-responders, sample size, normalization metric, and role in the analysis. The main dataset is GSE164458 with the highest sample size, used for the primary task. The independent validation dataset is GSE163882, and the auxiliary datasets are GSE194040, GSE41998, and GSE123845.**(B)** AUROC curves for TNBC-TAC independent validation (GSE123845) using auxiliary and main datasets (GSE163882). The plot shows the performance of different learning methods, with all auxiliary learning methods outperforming traditional machine learning models. The AUROC values for each method are detailed in the legend.

Next, we compared auxiliary learning against traditional machine learning with batch-corrected data. The independent validation results, shown in Figure 4-5B, indicate that GRADNORM achieved the highest AUROC (0.72), followed by ARML (0.65), OL AUX (0.62), and ADALOSS (0.61). In contrast, traditional machine learning models, even those trained on batch-corrected integrated datasets, did not exceed an AUROC of 0.6, demonstrating the superior performance of auxiliary learning. Interestingly, application of auxiliary learning to batch-corrected data did not lead to further performance improvements **(Table 4-1, lower section)**. These findings underscore the effectiveness of auxiliary learning in integrating diverse datasets for the same task, without the need for batch effect correction.

## Discussion

This study demonstrated that integrating auxiliary learning methods can significantly enhance predictive performance for TNBC treatment responses, especially in scenarios with limited data. The benchmarking analysis indicated that several auxiliary learning methods, such as ARML and OL AUX, achieved higher AUROC values in cross-validation, with GRADNORM showing the best independent validation results. However, ADALOSS appeared to underperform across various tests, suggesting variability in effectiveness among auxiliary learning methods.

Auxiliary learning methods outperformed baseline models primarily because they leveraged related auxiliary tasks, which provided additional relevant information to the main task of treatment prediction. This approach is particularly advantageous when sample sizes are small or when predicting responses to specific treatment arms, as the auxiliary tasks can help mitigate data scarcity by incorporating supplementary knowledge that enhances model learning.

However, there are limitations to this approach, including the risk of overfitting when the training and evaluation datasets contain very small or imbalanced labels. Future work should focus on expanding this approach by incorporating diverse omics data from clinical trials, which could enhance the model’s robustness and broaden its applicability. By applying these methods to omics data from TNBC patients, there is potential to develop a clinical tool that recommends the most effective treatment combinations, ultimately advancing personalized medicine for this challenging cancer subtype.

## Acknowledgments & Funding

B.Z. is a CPRIT Scholar in Cancer Research and a McNair Scholar. The content is solely the responsibility of the authors and does not necessarily represent the official views of the National Institutes of Health.

National Institute of General Medical Sciences [T32GM136554]; Cancer Prevention and Research Institute of Texas [RR160027 and RP220050]; The Robert and Janice McNair Foundation. Funding for open access charge: Cancer Prevention and Research Institute of Texas.

---

**Table 1.**
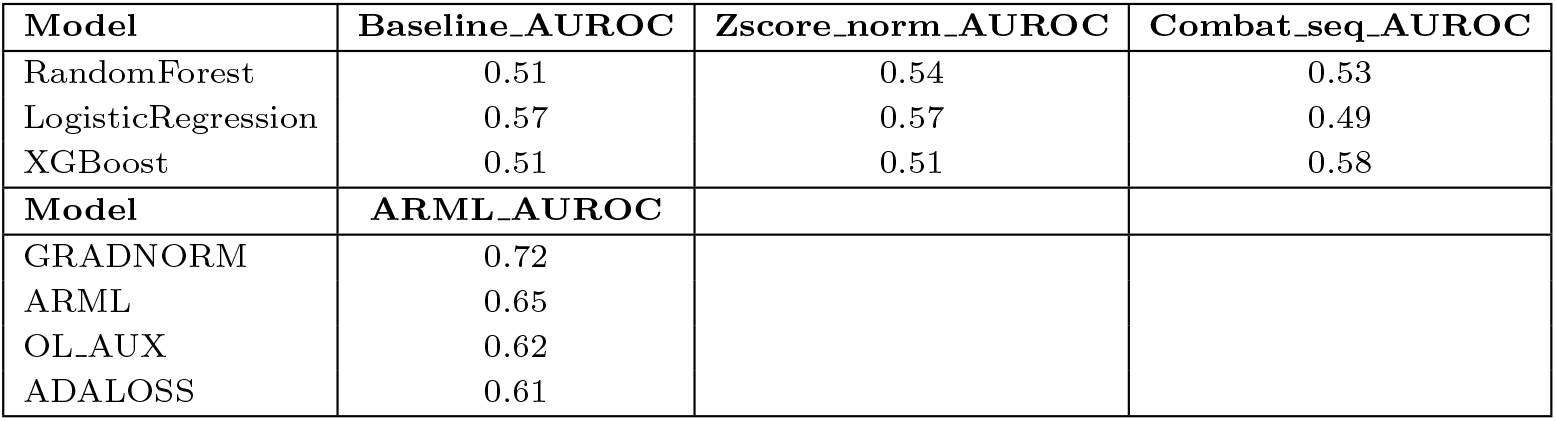
Comparison of AUROCs for GSE123845 independent validation results from supervised learning models (Random Forest, Logistic Regression, and XGBoost) using baseline, z-score normalization, and combat-seq normalization techniques. Additionally, the table presents the AUROCs for ARML.

**Supplementary Fig. 1.**
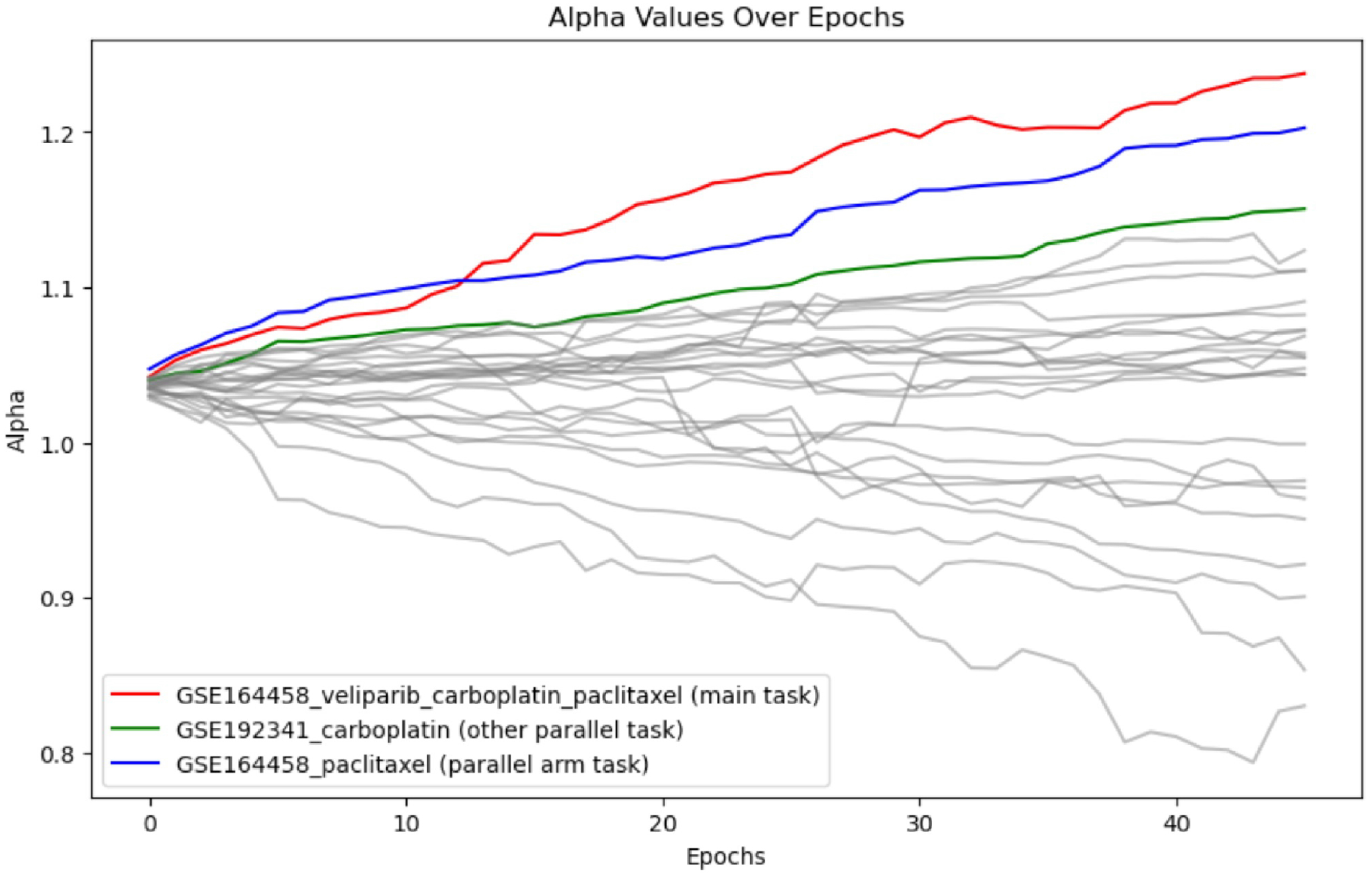
Alpha values over epochs for predicting outcomes for the GSE164458 veliparib carboplatin paclitaxel (main task). The plot shows alpha values for the main task (red line), and two parallel arm datasets: GSE192341 carboplatin (blue line) and GSE164458 paclitaxel (green line). The plot highlights that the parallel arm datasets, involving other platinum agents such as carboplatin and paclitaxel, significantly contribute to the model’s predictions, especially during the early epochs. The gray lines represent the individual trajectories of other tasks, showing varying degrees of contribution over time.

**Supplemental Table 1.** Comprehensive median 30 MCCV AUROCs across 27 TNBC treatment arms.

| Study | ARML | ADALOSS | GRADNORM | OL_AUX | XGBoost | RandomForest | LogisticRegression | MLP |
| --- | --- | --- | --- | --- | --- | --- | --- | --- |
| GSE123845_TAC | 0.3247 | 0.3563 | 0.232 | 0.2933 | 0.5495 | 0.5693 | 0.5029 | 0.5639 |
| GSE140494 | 0.6796 | 0.5986 | 0.6645 | 0.5408 | 0.5873 | 0.5113 | 0.5204 | 0.5253 |
| GSE163882 | 0.4455 | 0.4781 | 0.4793 | 0.4655 | 0.5006 | 0.5046 | 0.4902 | 0.5074 |
| GSE164458_carboplatin_paclitaxel | 0.5063 | 0.5373 | 0.5376 | 0.5061 | 0.4792 | 0.4884 | 0.5298 | 0.5571 |
| GSE164458_paclitaxel | 0.5933 | 0.5402 | 0.5279 | 0.5915 | 0.5442 | 0.5576 | 0.5583 | 0.5283 |
| GSE164458_veliparib_carboplatin_paclitaxel | 0.539 | 0.4931 | 0.5453 | 0.5244 | 0.5407 | 0.5344 | 0.5436 | 0.5232 |
| GSE16446 | 0.3634 | 0.4401 | 0.3105 | 0.2239 | 0.5895 | 0.5592 | 0.533 | 0.5333 |
| GSE192341 | 0.6148 | 0.5619 | 0.6035 | 0.6346 | 0.5212 | 0.535 | 0.3985 | 0.5325 |
| GSE194040_Paclitaxel | 0.5378 | 0.4368 | 0.481 | 0.4855 | 0.521 | 0.496 | 0.4693 | 0.5452 |
| GSE194040_Paclitaxel_ABT888_carboplatin | 0.6521 | 0.6386 | 0.7195 | 0.6226 | 0.7685 | 0.8048 | 0.7717 | 0.5761 |
| GSE194040_Paclitaxel_AMG386 | 0.4325 | 0.504 | 0.5006 | 0.4716 | 0.5272 | 0.5307 | 0.5675 | 0.5187 |
| GSE194040_Paclitaxel_Ganetespib | 0.578 | 0.5673 | 0.637 | 0.6293 | 0.7533 | 0.7204 | 0.6523 | 0.4505 |
| GSE194040_Paclitaxel_Ganitumab | 0.2866 | 0.3827 | 0.3181 | 0.3991 | 0.5556 | 0.5716 | 0.5731 | 0.5526 |
| GSE194040_Paclitaxel_MK_2206 | 0.5112 | 0.4315 | 0.5132 | 0.4693 | 0.4567 | 0.4682 | 0.48 | 0.4521 |
| GSE194040_Paclitaxel_Neratinib | 0.5187 | 0.5123 | 0.4656 | 0.5257 | 0.4302 | 0.433 | 0.493 | 0.4769 |
| GSE194040_Paclitaxel_Pembrolizumab | 0.6906 | 0.6366 | 0.6956 | 0.6744 | 0.5156 | 0.5273 | 0.5065 | 0.511 |
| GSE20194 | 0.448 | 0.4858 | 0.4803 | 0.414 | 0.5271 | 0.554 | 0.5545 | 0.5474 |
| GSE20271_FAC | 0.6832 | 0.6567 | 0.9018 | 0.5615 | 0.5496 | 0.5898 | 0.5724 | 0.5463 |
| GSE20271_TFAC | 0.5313 | 0.5354 | 0.5339 | 0.5879 | 0.49 | 0.5273 | 0.5298 | 0.5337 |
| GSE22093 | 0.398 | 0.3171 | 0.3766 | 0.4005 | 0.5896 | 0.528 | 0.6148 | 0.5635 |
| GSE23988 | 0.7788 | 0.6608 | 0.7636 | 0.92 | 0.7385 | 0.7421 | 0.7094 | 0.5722 |
| GSE25055 | 0.6313 | 0.5584 | 0.6379 | 0.5561 | 0.5339 | 0.752 | 0.5117 | 0.587 |
| GSE25065 | 0.3788 | 0.4477 | 0.4249 | 0.4815 | 0.5401 | 0.5321 | 0.5921 | 0.5229 |
| GSE41998_Ixabepilone | 0.4872 | 0.4943 | 0.474 | 0.4495 | 0.5248 | 0.5506 | 0.516 | 0.5109 |
| GSE41998_Paclitaxel | 0.6327 | 0.554 | 0.6075 | 0.5977 | 0.5543 | 0.56 | 0.5131 | 0.6049 |
| GSE42822_FEC_TX | 0.4843 | 0.6095 | 0.5311 | 0.6255 | 0.5451 | 0.5848 | 0.5826 | 0.5828 |
| GSE4779 | 0.555 | 0.4989 | 0.5935 | 0.5897 | 0.5575 | 0.5221 | 0.5525 | 0.563 |

