## Supplementary figures and images for "Optimizing Neoadjuvant Treatment Response Prediction for Triple-Negative Breast Cancer Using Clinical Trial Data and Deep Auxiliary Learning"

### Supplemental Figure 1

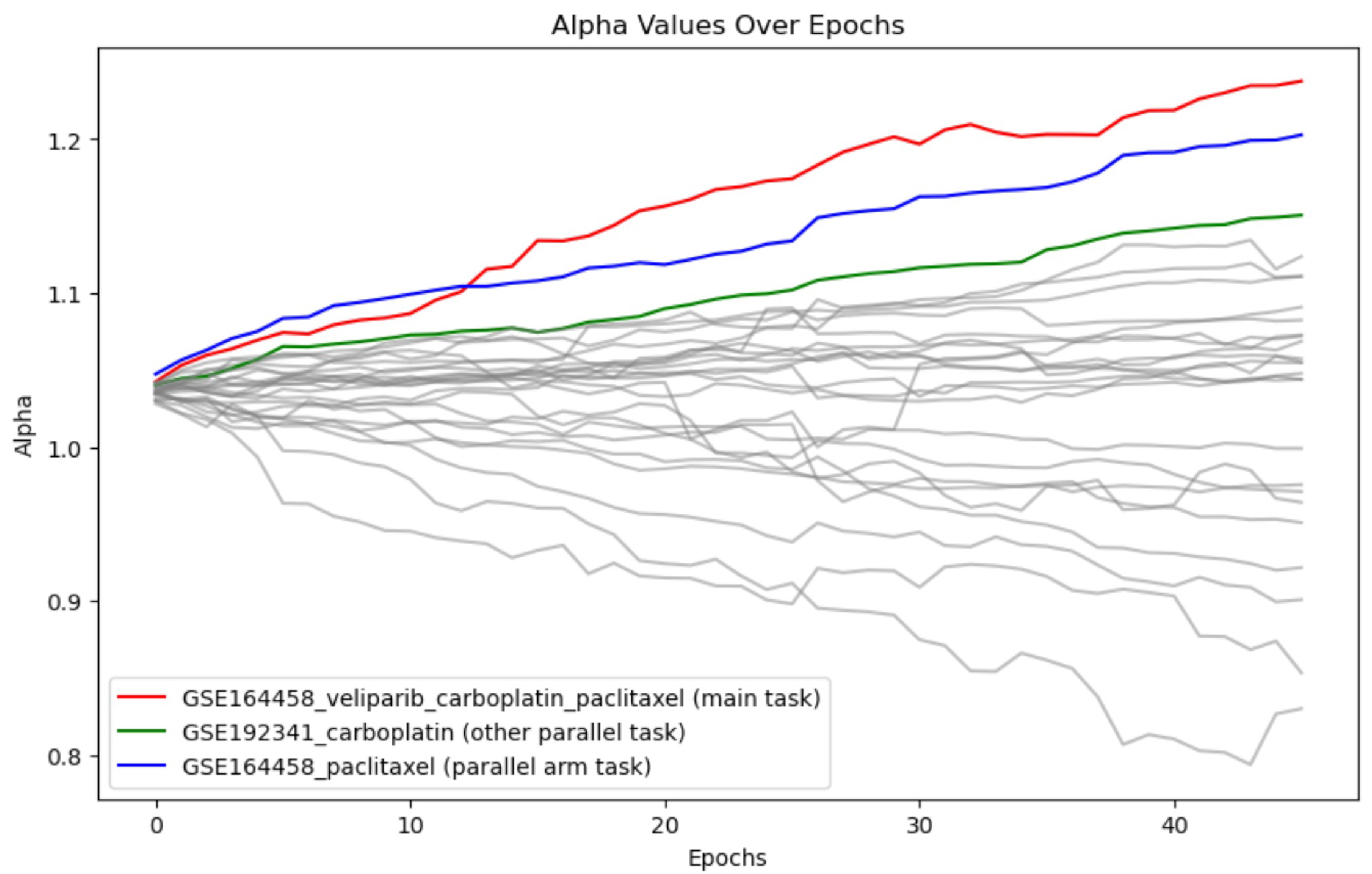
